# Natural resistance to worms exacerbates bovine tuberculosis severity independently of worm coinfection

**DOI:** 10.1101/2020.08.26.262683

**Authors:** Vanessa O. Ezenwa, Sarah A. Budischak, Peter Buss, Mauricio Seguel, Gordon Luikart, Anna E. Jolles, Kaori Sakamoto

## Abstract

Pathogen interactions arising during coinfection can exacerbate disease severity, for example, when the immune response mounted against one pathogen negatively affects defense of another. It is also possible that host immune responses to a pathogen shaped by historical evolutionary interactions between host and pathogen, may modify host immune defenses in ways that have repercussions for other pathogens. In this case, negative interactions between two pathogens could emerge even in the absence of concurrent infection. Parasitic worms and tuberculosis (TB) are involved in one of the most geographically extensive of pathogen interactions, and during coinfection, worms can exacerbate TB disease outcomes. Here, we show that in a wild mammal, natural resistance to worms affects bovine tuberculosis (BTB) severity independently of active worm infection. We found that worm-resistant individuals were more likely to die of BTB than were non-resistant individuals, and their disease progressed more quickly. Anthelmintic treatment moderated, but did not eliminate, the resistance effect, and the effects of resistance and treatment were additive with untreated, resistant individuals experiencing the highest mortality. Interestingly, resistance and anthelmintic treatment had non-overlapping effects on BTB pathology. The effects of resistance manifested in the lungs (the primary site of BTB infection), while the effects of treatment manifested almost entirely in the lymph nodes (the site of disseminated disease), suggesting that resistance and active worm infection affect BTB progression via distinct mechanisms. Our findings reveal that interactions between pathogens can occur as a consequence of processes arising on very different timescales.

## Introduction

Interactions between pathogens co-occurring within a single host can have profound effects on infection outcomes, ranging from the severity of clinical disease in individual hosts to the rate of disease spread across populations (1-3). Because most hosts are commonly infected by more than one type of pathogen at a time (4), understanding the consequences of pathogen interactions during concurrent infection (or coinfection) is essential for effective disease management and control. While many studies focus on pathogen interactions that are the result of one pathogen responding to the simultaneous presence of another (5), two pathogens need not overlap in time to interact with one another. For example, heterologous immunity, where prior exposure or infection with one pathogen modifies the immune response to another, can drive both positive and negative interactions between pathogens (6). This phenomenon highlights how modifications of the host immune system by one pathogen that occur during the lifetime of a host (i.e. in ecological time) can shape future responses to secondary pathogens. Likewise, strong selection pressure imposed by pathogens on hosts, particularly on immune function (7), can result in modifications of the host immune system that occur over generations (*i*.*e*. in evolutionary time), a process which should also affect responses to secondary infections. In this case, a historical population-level response to selection by one pathogen may generate heritable differences among individuals in contemporary responses to another. Crucially, ecological- *vs*. evolutionary-scale interactions between pathogens may operate for different reasons, so distinguishing between the two is integral to understanding both the mechanistic basis and consequences of these interactions.

Helminths, or parasitic worms, and tuberculosis (TB) are involved in one of the most geographically extensive of pathogen interactions (2, 8). Both pathogens affect approximately one-third of the world’s human population and are widespread in domestic and wild animals (9-11). Caused by bacteria in the *Mycobacterium tuberculosis* complex, including *M. tuberculosis* (*Mtb*) the causative agent of human tuberculosis and *M. bovis* (*Mb*) the causative agent of bovine tuberculosis, TB is responsible for 2 million human deaths (12) and 25% of all disease-related cattle deaths (13), annually. In humans, about 10% of individuals infected with *Mtb* progress to active pulmonary disease, but the mechanisms underlying progression to active TB are poorly defined (14). Accumulating evidence suggests that coinfection with worms may be a factor in TB disease progression (15, 16), although some studies do not support this link, highlighting the complex nature of worm-TB interactions (17). Interestingly, research in lab animals suggests that enhanced immunity (i.e. resistance) to worms can compromise a host’s ability to control TB even in the absence of active worm infection (18-21), implying that evolved defenses against worms may independently affect the response to TB. Considered in light of widespread worm resistance in human and animal populations (22, 23) and the broad geographic coincidence of worms and TB, worm-TB interactions may represent an illustrative case where variation in evolved resistance to one pathogen (worms) contributes to variable responses to another (TB).

In this study, we tested the hypothesis that resistance to worms modifies the host response to TB. To do this, we monitored gastrointestinal worm (specifically strongyle nematode) and *Mb* infections in a cohort of wild African buffalo (*Syncerus caffer*) to assess the effects of natural variation in worm resistance on the incidence, severity, and progression of bovine tuberculosis (BTB). In previous work, we demonstrated the presence of an ecological interaction between worms and BTB in buffalo by using showing that clearance of active worm infection via anthelmintic treatment reduces BTB-associated mortality (24). Thus, we took advantage of the fact that half of our study animals were subject to long-term deworming, to compare the relative effects of worm coinfection versus natural worm resistance on BTB outcomes. We found evidence of a genetic basis to worm resistance in buffalo and that buffalo with resistance to worms were more severely affected by BTB in terms of both mortality risk and disease progression. However, the mechanisms by which natural variation in the host response to worms was associated with BTB progression appeared to be distinct from the effects of anthelmintic treatment. Our results suggest that negative effects of worms on BTB outcomes occur as a result of both concurrent worm infection and genetically-based differences in host responsiveness to worms. This discovery fundamentally alters our understanding of the timescales over which worms and TB interact in real-world populations.

## Results

We tracked 209 female African buffalo over four years in Kruger National Park, South Africa. One hundred and twelve animals (53%) were infected with worms at initial sampling, and the number of worm eggs shed in the feces of all individuals (hereafter referred to as the fecal egg count [FEC]) varied from 0-850 eggs per gram of feces (Mean ± SE = 96.9 ±10). At the first and all subsequent sampling events, approximately half of the study animals, designated as the treatment group, received an anthelmintic drug to control their worm infections. This treatment regime effectively reduced worm egg shedding (24), a reliable proxy for adult worm burden in this study population (25). Of the subset of control animals that did not receive anthelmintic treatment, FEC was significantly repeatable within an individual over time (R [control animals, n = 103 individuals, 651 observations, mean [range] = 6 [2-9] observations per individual] = 0.41, 95% CI: 0.31-0.49; *P* < 0.0001), revealing a high degree of natural within-individual consistency in the magnitude of worm egg shedding. FEC is a widely-used proxy of worm resistance in livestock (26), so to investigate the potential source of this consistent variation in egg shedding in buffalo we classified animals into discrete categories based on egg shedding levels at initial sampling. We used a model-based approach to identify clusters in the FEC data and individuals were classified into two groups based on these clusters: ‘low’ FEC (<100) representing the largest cluster and ‘high’ FEC (≥100) representing all other clusters (*SI Appendix*, Fig. S1). We then tested whether this ‘low’ vs. ‘high’ FEC phenotype was predictive of variation in the host immune response to worm infection and looked for evidence of a genetic basis to the phenotype. The approach of discretizing FEC values allowed us to identify patterns that might have been obscured due to the highly skewed distribution of the FEC dataset (Fig. 1A) and potential non-linear relationships between FEC and TB disease progression.

**Figure 1.**
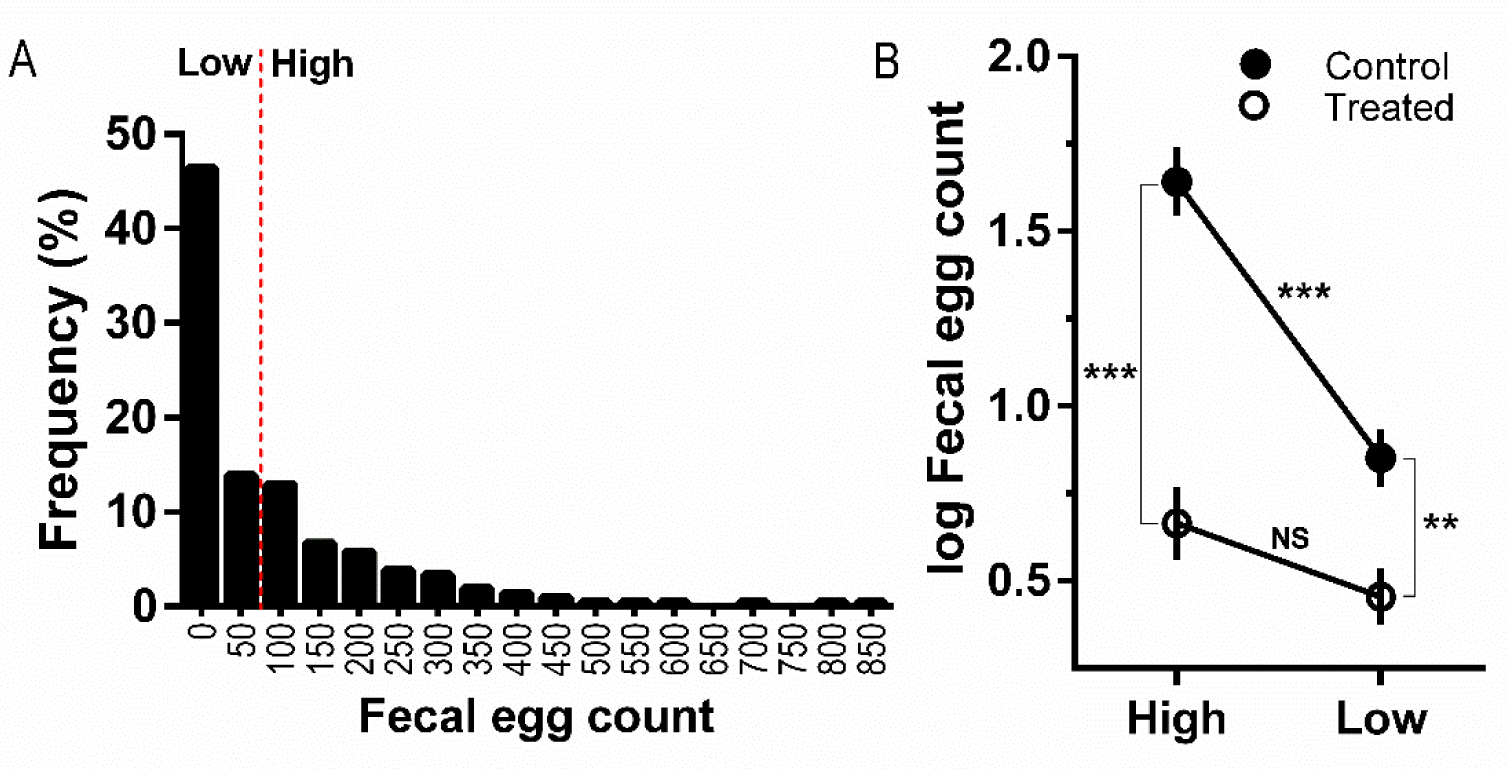
Fecal egg counts (FEC) at initial capture were highly aggregated across individuals, and worm egg shedding phenotype (high or low) at first capture interacted with anthelmintic treatment to determine shedding throughout the study. (A) At the beginning of the study, 60% of buffalo were shedding less than 100 worm eggs per gram feces and were classified as the ‘low’ phenotype, while 40% were shedding greater than 100 eggs per gram feces and were classified as the ‘high’ phenotype (*SI Appendix*, Fig. S1). (B) Throughout the study, high FEC individuals in the control group (control-high) shed significantly more worm eggs than either low FEC individuals in the control group (control-low) or high FEC individuals in the treated group (treated-high). The magnitude of the effect of worm phenotype on worm egg shedding (control-high – control low: 0.79, *SI Appendix*, Table S2) was nearly equivalent to the effect of anthelmintic treatment (control-high – treated-high: 0.98, *SI Appendix*, Table S2). Fecal egg counts were log (x+1) transformed, symbols show means and standard errors, asterisks denote significant differences (***< 0.001, ** < 0.01, NS = not significant).

Sixty percent of animals fell into the low FEC category and 40% fell into the high FEC category (Fig. 1A), and the probability of an individual expressing a low *vs*. high FEC phenotype was repeatable over time (R logit-link approximation [control animals, n = 103 individuals, 651 observations] = 0.999, 95% CI: 0.995-0.999; *P* < 0.0001), indicating that our discretized measure of FEC phenotype captured the consistent variation in egg shedding apparent in the continuous FEC metric. Furthermore, FEC phenotype at initial capture was associated with differences in worm egg shedding throughout the study nearly equivalent in magnitude to the effect of anthelmintic treatment (*SI Appendix*, Table S1-S2). Accounting for a range of factors associated with worm exposure risk (age, herd, season), we found that among controls, individuals with the high phenotype shed significantly more worm eggs than individuals with the low phenotype, but treatment eliminated the difference between high and low individuals (Fig. 1B, *SI Appendix*, Table S2). Moreover, the difference in egg shedding between control-high and control-low individuals was similar to the difference between control-high and treated-high individuals (Fig. 1B, *SI Appendix*, Table S2), indicating that FEC phenotype had effects on worm egg shedding almost as potent as anthelmintic treatment. Crucially, the high-low phenotype was reflective of differences in key immune responses to worms. Among control animals, low individuals had more eosinophils, mast cells, and IgA at the primary sites of worm infection (abomasum and small intestine) compared to high individuals (Wilcoxon rank sum test [control animals only]: abomasum (n = 7 low, 7 high, eosinophils (Fig. 2A): *Z* = -2.41, *P* = 0.0159; mast cells (Fig. 2B): *Z* = -2.13, *P* = 0.0328; IgA (Fig. 2C): *Z* = -3.07, *P* = 0.00006; intestine (n = 6 low, 6 high), eosinophils: *Z* = -1.86, *P* = 0.0632; mast cells: *Z* = -2.81, *P* = 0.0049; IgA: *Z* = -2.80, *P* = 0.0021). The fact that all three of these well-known markers of anti-helminth immunity (27) were higher in low FEC individuals supports the presence of a more robust worm resistance response in these animals.

**Figure 2.**
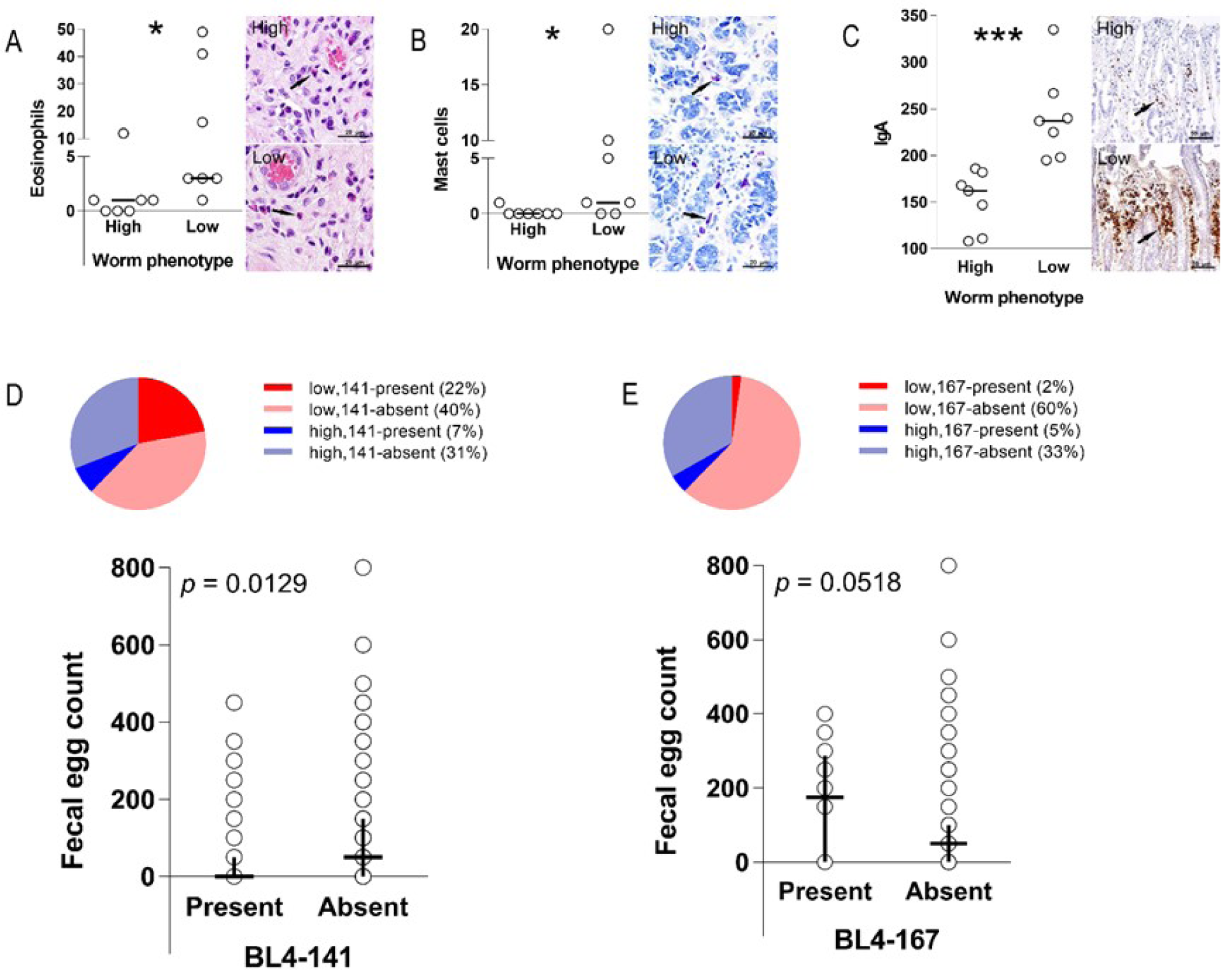
The high-low fecal egg count (FEC) phenotype in buffalo reflects variation in the immune response to worms at the site of infection in the gut and has an underlying genetic basis. Low FEC individuals had significantly more (A) eosinophils (cells per 2mm^2^), (B) mast cells (cells per 2mm^2^), and (C) IgA (number IgA+ leukocytes) in the abomasum than high FEC individuals. Horizontal bars show median values, open circles are raw data points, and asterisks denote significant differences (***< 0.001, * < 0.05). Furthermore, FEC phenotype was associated with polymorphism and the BL4 locus near the interferon gamma gene. (D) Allele BL4-141 was over-represented among low FEC (dark red *vs*. light red) compared to high FEC (dark blue *vs*. light blue) individuals, and individuals carrying allele 141 shed significantly fewer worm eggs (cross represents median and interquartile range, open circles show full range of FEC values). (E) In contrast, allele BL4-167 was over-represented among high FEC (dark blue *vs*. light blue), compared to low FEC individuals (dark red *vs*. light red), and individuals carrying this allele tended to shed more worm eggs (cross represents median and interquartile range, open circles show full range of FEC values).

FEC phenotype was also associated with allelic variation near the interferon gamma gene (IFNG), a gene region that has previously been linked to variation in nematode egg counts in domestic sheep (28), feral Soay sheep (29), and buffalo (30). Focusing on the BL4 locus, which is located 3.6cM upstream of IFNG (31), we identified nine alleles, two of which, BL4-141 and BL4-167, were associated with FEC phenotype. BL4-141 occurred at a frequency of 29% in the study population (51 out of 177 sampled individuals), and low animals were significantly more likely to carry this allele (high = 12/67, low = 39/110, Pearson’s chi-squared test: χ^2^ = 6.25, *P* = 0.0124; Fig. 2D). By contrast, BL4-167 was relatively rare in the study population, occurring at a frequency of only 7%, and in this case, high animals were significantly more likely to be carriers of the allele (high = 8/67, low = 4/110, Pearson’s chi-squared test: χ^2^ = 4.54, *P* = 0.033; Fig. 2E). Further supporting these patterns, the presence of both alleles was correlated with continuous FEC at initial sampling. Individuals carrying BL4-141 were shedding significantly fewer worm eggs (Wilcoxon rank sum test: *Z* = -2.49, *P* = 0.0129; Fig. 1D), while those carrying BL4-167 tended to shed more eggs (*Z* = 1.94, *P* = 0.0518; Fig. 2E). In combination, both the immunological and genetic patterns suggest that the high-low FEC phenotype in buffalo is a reliable proxy for genetically-based differences in natural resistance to worm infection.

When we examined the consequences of FEC phenotype (herafter called ‘worm resistance’ phenotype) for BTB disease, we found that resistance did not affect the risk of buffalo acquiring BTB (high = 29/77, low = 37/118; Hazard Ratio [HR] = 0.708, 95% CI: 0.427-1.177, *P* = 0.341, *SI Appendix*, Table S3), but was a strong predictor of mortality risk after BTB infection. Among BTB-infected individuals, mortality was approximately 6-times higher among low compared to high individuals (high = 2/24, low = 10/32; HR = 5.68, 95% CI: 1.08-29.9, *P* = 0.0089, Fig. 3, *SI Appendix*, Table S4). This worm resistance effect was independent of anthelmintic treatment status, but lack of treatment had an additive effect on mortality risk (*SI Appendix*, Table S3). There was a 26% cumulative mortality difference between control-high and control-low individuals (Fig. 3). However, cumulative mortality of treated-high individuals was 0% compared to 14%% in treated-low individuals (Fig. 3), suggesting that treatment dampened the effect of worm resistance phenotype although worm resistance still imposed a BTB-associated mortality cost in spite of significantly reduced worm burdens due to treatment. Treatment reduced BTB-associated mortality in both high and low individuals, as reflected by 18% and 30% reductions in cumulative mortality in control *vs*. treated, high and low individuals, respectively (Fig. 3). And, the additive negative effect of worm resistance and lack of treatment was most apparent in the substantial cumulative mortality differential between treated-high and control-low individuals (0% vs. 44%; Fig. 3). Importantly, there was no effect of worm resistance phenotype on mortality among individuals that were not infected with BTB (*SI Appendix*, Table S5), implicating an interaction between worm resistance phenotype and BTB as the driver of the resistance-based mortality pattern. Finally, when we examined associations between continous FEC and mortality, we found patterns consistent with results based on worm resistance phenotype. Among BTB-infected individuals, those with lower FECs showed a tendency towards higher mortality (*P* = 0.07, *SI Appendix*, Table S6), whereas this trend disappeared among individuals not infected with BTB (*P* = 0.42, *SI Appendix*, Table S7). These patterns corroborate that our discretized FEC (i.e. worm resistance) phenotype captures an important underlying biological phenomenon.

**Figure 3.**
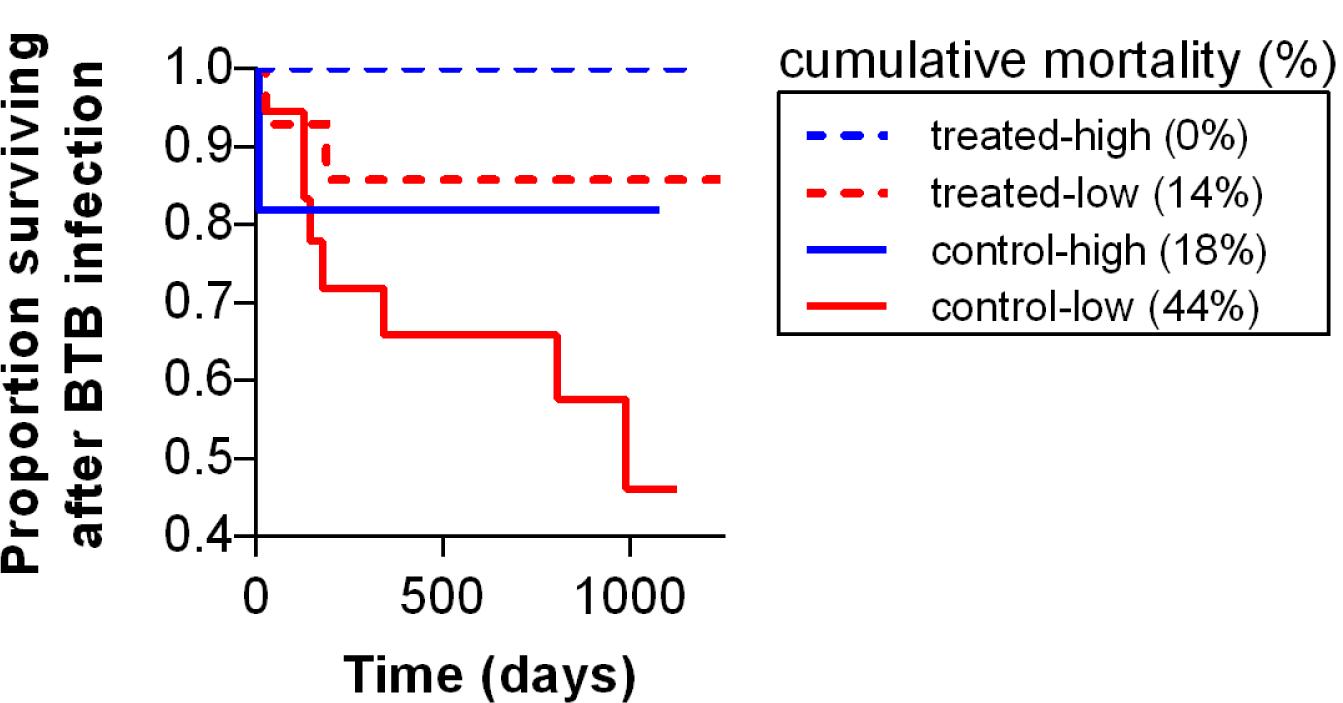
Worm resistance phenotype affects the survival of buffalo infected with BTB, and anthelmintic treatment reduces, but does not eliminate this effect. Survival curves showing the proportion of worm resistant (low worm fecal egg count [FEC] phenotype) and non-resistant (high FEC phenotype) buffalo by anthelmintic-treatment and control status surviving BTB infection as a function of time in days. The probability of death given BTB infection was significantly higher for low FEC individuals compared to high FEC individuals and worm resistance effects were independent of anthelmintic treatment (*SI Appendix*, Table S4). Cumulative mortality rates across all groups show that while treatment moderated the mortality effect of worm resistance (treated-high *vs*. control-high and treated-low *vs*. control-low), resistant animals treated for worms (treated-low) still suffered a mortality cost of resistance.

To better understand how worm resistance phenotype and anthelmintic treatment accelerated BTB mortality, we performed gross and histolopathological examination of the lungs and respiratory lymph nodes of a subset of BTB-infected individuals to identify the effect of these traits on BTB disease progression. The lungs are the primary site of BTB infection and the lymph nodes, a common site of extrapulmonary infection, reflect dissemination of the disease (32). In the lung, we quantified the host’s ability to control bacterial infection (number of lesions, multinucleated giant cell [MNGC] formation), the length and severity of infection (lesion stage), and the degree of tissue damage within lesions (presence of mineralization), including the inability to return to normal tissue architecture (fibrosis). After accounting for the duration of BTB infection, subjects with the low phenotype tended to have more lung lesions (Table 1). Lesions in this group of animals were also significantly more likely to have MNGCs present (Table 1 and *SI Appendix*, Fig. S2A) and were at a significantly more advanced stage (Table 1), both of which are indicators of less effectively controlled BTB infection. Only one index of disease in the lung, the presence of fibrosis within lesions, was affected by anthelmintic treatment and not worm resistance phenotype. Treated animals had a higher degree of fibrosis within lung lesions (Table 1), which suggests that treatment was associated with an inability to reverse tissue damage in the lung, while worm infection promoted tissue regeneration.

**Table 1.**
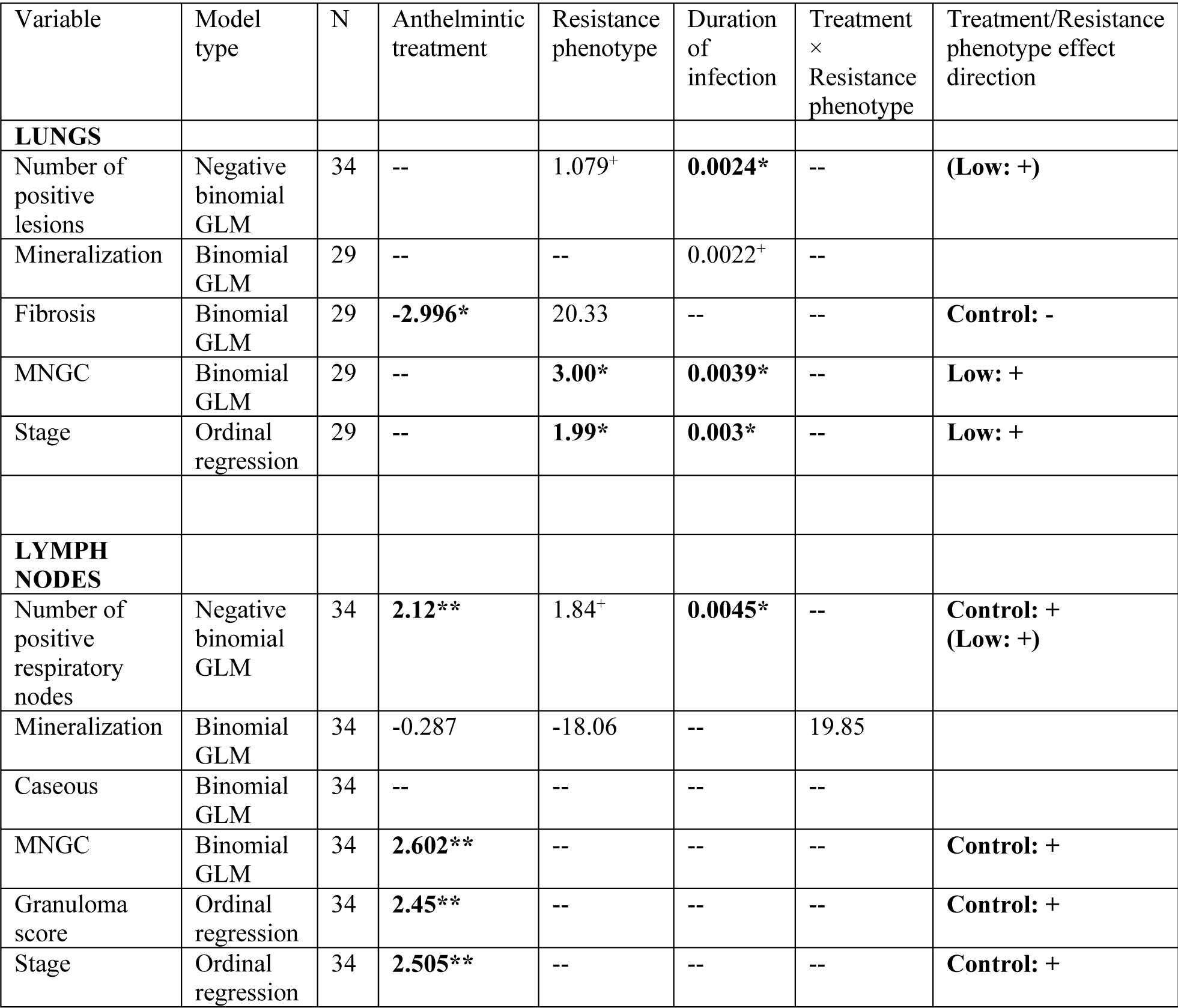
Effects of anthelmintic treatment, worm resistance phenotype, and duration of infection on BTB severity quantified by gross and histolopathological examination of the lungs and respiratory lymph nodes. In the lungs, severity was measured as the number of BTB lesions and multinucleated giant cell [MNGC] formation (indicators of the host’s ability to control bacterial infection), lesion stage (indicator of the length and severity of infection), and the presence of mineralization and fibrosis (indicators of the degree of tissue damage within lesions). In the lymph nodes, severity was measured as the number of respiratory nodes with BTB lesions (indicator of disease dissemination), the retropharyngeal lymph node (RPLN) granuloma score and stage (indicators of the distribution and severity of infection), and mineralization, caseous necrosis and MNGC formation (indicators of the degree of tissue damage within RPLN lesions). Full models of the structure: Y ∼ anthelmintic treatment + resistance phenotype + infection duration + treatment × resistance phenotype were run for each variable and final models were selected using AIC-based model simplification. Values represent estimates from the final models. Significant factors appear in bold (significance codes: ***< 0.001, ** < 0.01, * < 0.05, ^+^ < 0.1). Effect description summarizes the direction of each significant treatment and resistance phenotype effect; trends appear in parenthesis.

| Variable | Model type | N | Anthelmintic treatment | Resistance phenotype | Duration of infection | Treatment × Resistance phenotype | Treatment/Resistance phenotype effect direction |
| --- | --- | --- | --- | --- | --- | --- | --- |
| **LUNGS** |  |  |  |  |  |  |  |
| Number of positive lesions | Negative binomial GLM | 34 | -- | 1.079<sup>+</sup> | **0.0024\*** | -- | **(Low: +)** |
| Mineralization | Binomial GLM | 29 | -- | -- | 0.0022<sup>+</sup> | -- |  |
| Fibrosis | Binomial GLM | 29 | **-2.996\*** | 20.33 | -- | -- | **Control: -** |
| MNGC | Binomial GLM | 29 | -- | **3.00\*** | **0.0039\*** | -- | **Low: +** |
| Stage | Ordinal regression | 29 | -- | **1.99\*** | **0.003\*** | -- | **Low: +** |
| **LYMPH NODES** |  |  |  |  |  |  |  |
| Number of positive respiratory nodes | Negative binomial GLM | 34 | **2.12\*\*** | 1.84<sup>+</sup> | **0.0045\*** | -- | **Control: + (Low: +)** |
| Mineralization | Binomial GLM | 34 | -0.287 | -18.06 | -- | 19.85 |  |
| Caseous | Binomial GLM | 34 | -- | -- | -- | -- |  |
| MNGC | Binomial GLM | 34 | **2.602\*\*** | -- | -- | -- | **Control: +** |
| Granuloma score | Ordinal regression | 34 | **2.45\*\*** | -- | -- | -- | **Control: +** |
| Stage | Ordinal regression | 34 | **2.505\*\*** | -- | -- | -- | **Control: +** |

In contrast to the lungs, BTB progression in the lymph nodes was exclusively associated with anthelmintic treatment status and not worm resistance phenotype (Table 1). To evaluate lymph node disease, we quantified the number of respiratory lymph nodes with BTB lesions and also assessed lesion severity within the retropharyngeal lymph node (RPLN). In the RPLN, we quantified the distribution (granuloma score), severity (stage), and degree of tissue damage (mineralization, caseous necrosis, and MNGC formation). Animals treated with an anthelmintic had signficantly fewer positive lymph nodes (Table 1), and lesions in the RPLN were less advanced in terms of disease stage, with less node involvement (i.e. lower granuloma score) and fewer MNGCs (Table 1 and *SI Appendix*, Fig. S2B).

## Discussion

Our study revealed that resistance to worms, like worm coinfection, affects the host response to BTB infection. We also showed that worm resistance and anthelmintic treatment had distinct effects on BTB disease progression. Our observation that TB severity is independently affected by the timescale of the host response to worms (*i*.*e*. ecological [active infection] *vs*. evolutionary [resistance]) has profound implications for understanding variation in TB outcomes in natural populations. Specifically, our finding that worm-resistant individuals infected with BTB suffered from more severe disease progression and higher mortality, even when they were treated for worms, suggests that real-time interactions between worms and TB are not required for resistance-induced negative TB outcomes to manifest. Rather, genetic variation in worm resistance, likely driven by historical evolutionary interactions between hosts and worms (33, 34), is sufficient to generate variation in TB disease progression. Counterintuitively, this means that hosts who control worms best and have the lowest rates of worm coinfection may be as compromised in their ability to control TB as hosts with high worm burdens.

In livestock, fecal egg counts (FECs) are a widely used as a proxy for worm resistance (35). In sheep, for example, FECs are a heritable trait predictive of variation in anti-helminth immunity (36). We showed that that FEC phenotype in buffalo was repeatable, linked to differences in anti-helminth immunity, and associated with variation at an immune gene known to play a role in worm resistance. In aggregate, this body of evidence supports the use of FEC phenotype as a proxy for worm resistance in our population. Crucially, our proxy for worm resistance was highly predictive of variation in mortality among BTB-infected, but not uninfected, buffalo, suggesting that worm resistance phenotype and BTB interacted to drive mortality outcomes. Furthermore, the effect of worm resistance on BTB mortality was independent of the presence of worms, and the dual effects of worm resistance and worm presence were additive, such that individuals who were both resistant to worms and actively infected with worms experienced the most severe mortality costs. The additive effect implies that worm-TB interactions can arise from distinct processes.

The non-overlapping effects of resistance and anthelmintic treatment on BTB pathology support the idea that treatment and resistance affect BTB progression via distinct mechanisms. We found that the effects of worm resistance on BTB pathology manifested primarily in the lungs, whereas the effects of anthelmintic treatment manifested primarily in the lymph nodes. The most commonly cited mechanism by which worms affect TB is via modulation of the host immune response. Specifically, activation of T helper cell 2 (Th2) immune responses by worms and concomitant downregulation of antibacterial T helper cell 1 (Th1) responses has been identified as a central mechanism linking worm infection to increased TB severity (37-40). More recently, induction of alternatively activated macrophages (AAMs) in the lungs during worm coinfection has been associated with compromised TB control (18, 19). Intriguingly, AAMs are also associated with enhanced immunity to worms (20, 21), and genetic variability in arginase-1 activity, the key enzyme expressed by AAMs, affects the outcome of *Mtb* infection even in the absence of worm infection (18, 41). Taken together, these observations suggest a potential mechanistic pathway (i.e. level of AAM activation) by which resistance to worms may independently affect TB control and disease progression.

The idea that AAM activation may explain the link we found between worm resistance and BTB disease progression is indirectly supported by several of our histopathology observations. First, MNGC formation is typically associated with chronic inflammation in the lung (42), and the more severe lung inflammatory state we described in worm resistant, BTB-infected animals (Fig. S2A) mirrors observations that *Mtb*-infected mice with increased arginase-1 activity show greater inflammatory damage in the lungs (18). Second, an AAM-mediated inflammatory effect would explain why we saw only a weak association between resistance phenotype and the number of lung lesions, since the primary effect of worm resistance should operate via inflammation-induced pathology rather than via bacterial control. Indeed, if this hypothesis is correct, the lack of an effect of worm resistance phenotype on BTB progression in the lymph nodes may be a direct consequence of a less immunoreactive environment in the lymph node (explaining no effect on tissue architecture) and limited impact on bacterial replication (explaining the weak effect on bacterial dissemination to lymph nodes). Third, in direct contrast, the strong effect of anthelmintic treatment on the lymph nodes, particularly the reduction in numbers of *Mb-*infected nodes, is consistent with a dampening of worm-induced Th1 impairment by anthelmintic treatment and concomitant increase in host control over bacterial replication. Together, these observations suggest an intriguing role for AAM activation in explaining effects of worm resistance on BTB, however, future studies at the mechanistic level are required to test this hypothesis and the idea that distinct mechanisms underlie the effects of worm reistsance *vs*. anthelmintic treatment on BTB.

Irrespective of the precise mechanisms governing the effects of worm resistance and anthelmintic treatment on BTB disease progression, our results shed important new light on the nature and complexity of worm-TB interactions, specifically, and pathogen interactions more generally. Despite overwhelming laboratory evidence that worms and various microbes interact in ways that can be detrimental to hosts (2, 43), there is very little consensus on the consequences of these interactions for disease outcomes in real-world populations. Our work in wild buffalo reveals that interactions between the same pair of pathogens can originate from processes operating on very different timescales, driven by potentially distinct underlying mechanisms. This insight may be central to fully understanding of the dynamics of coinfection and implications for disease control. For TB, accounting for worm-TB interactions driven by processes operating on an evolutionary timescale might help reconcile inconsistencies in empirical patterns (*e*.*g*. 16) currently viewed solely through the lens of ecological time. There are also ramifications for disease control. For example, in humans, accounting for patterns of genetic variation in worm resistance may provide new insight in the search for biomarkers of TB disease progression. While in livestock, where breeding for worm resistance is an increasingly necessary alternative to anthelmintic drug use (26), it may be economically expedient to understand whether breeding for resistance is associated with unanticipated BTB-associated health costs.

## Materials and Methods

### Animal captures

Female African buffalo (*Syncerus caffer*) were captured in the southern portion of Kruger National Park (KNP), South Africa between June 2008 and August 2012. Animals were captured by helicopter in 2008, fitted with radio-collars, and then re-captured twice a year by vehicle. Animals were captured from two herds in different locations, the Crocodile Bridge (CB) herd and the adjacent Lower Sabie (LS) herd. At initial capture, we randomly assigned animals to an anthelmintic treatment group (fenbendazole bolus, Intervet) or control group. Individuals lost due to death or emigration during the study were replaced and replacements were assigned to the same experimental group as the original animal. Details on the efficacy of the anthelmintic treatment are described in (24). Animals were captured on average 6 times (range: 2-9 times), and at all captures, we collected fecal and blood samples to monitor worm infection and BTB infection status, respectively. We also assessed each animal’s age as described in (44). In total, we followed 209 unique individuals who were bovine tuberculosis (BTB)-negative at the beginning of the study and for whom data were available on worm infection at initial capture.

### Worm infection, resistance, and immunity

The gastrointestinal (GI) worm community of buffalo in our study was comprised of at least seven species of nematodes, dominated by the strongyle nematodes: *Cooperia fuelleborni, Haemonchus placei* and *Haemonchus bedfordi* (25). We monitored strongyle worm infection status using fecal samples collected at capture. Feces were collected directly from the rectum of immobilized animals, stored at 4°C in the field, and processed the same day. We quantified nematode egg output using a modification of the McMaster fecal egg counting technique with a sensitivity of 50 eggs per gram of feces (45). Based on the number of worm eggs individual’s shed at their initial capture, we classified animals into two categories (i.e. worm resistance phenotypes), ‘low’ versus ‘high’ fecal egg counts (FEC) using an unsupervised machine learning approach. We used a Gaussian mixture model implemented in the *mclust v5* package in R version 3.4.3 (46) to identify clusters in the FEC data. Model parameters were estimated using the Expectation-Maximization (EM) algorithm. The optimal model identified had 7 clusters (log-likelihood = -1221.863, n = 209, df = 14, BIC = -2518.518), with 126 of 209 samples placed in the first cluster and 83 samples placed in all remaining clusters (*SI Appendix*, Fig. S1). We used the cutoff value for cluster 1 (<100 FEC) to classify individuals into ‘low’ and ‘high’ FEC categories, respectively. Finally, we verified that these resistance phenotypes were associated with differences in immunological resistance by testing for an association between phenotype and key mucosal immunological responses to worm infection: numbers of eosinophils, mast cells, and total IgA secretion in the GI tract (27, 47).

We quantified GI tract immune responses to worms focusing on the abomasum and small intenstine (jejunum), the two primary sites of infection for our focal nematode species. GI tract tissue was collected from a subset of study animals euthanized and necropsied at the conclusion of the study. Euthanasia was performed as part of the KNP BTB surveillance program and implemented according to KNP standard operating procedure. The number of eosinophils was recorded by examining standard formalin-fixed, paraffin-embedded sections of abomasum and proximal jejunum tissue sections (8mm thick) stained with hematoxylin and eosin. For mast cells, replicates of the same tissue sections were stained with Giemsa stain and then counted. The presence of eosinoophilic granules and a polymorphic nucleus allowed the identification of eosinophils and the presence metachromatic granules facilitated the identification of mast cells (47). Eosinophil and mast cell counts are reported as the number of cells per 10 random high power fields at the villi lamina propria, which is equivalent to the number of cells per 2mm^2^.

IgA production was measured using immunohistochemistry. In this case, formalin-fixed, paraffin-embedded sections of abomasum and proximal jejeunum were deparafinized, rehydrated, and incubated with a diluted (1:800), rabbit, anti-bovine IgA primary antibody for 1 hour (Bethyl Laboratories, CO, USA, cat# A10-108A). After 30 minutes of incubation with biotinylated universal goat link (Dako, Carpinteria, CA, USA), visualization of antigen-antibody complexes was obtained via a 10-min incubation with streptavidin-conjugated horse radish peroxidase (Biocare, Chicago, IL, USA), followed by a 5 minute incubation with diaminobenzidine-peroxidase (Vector Laboratories, Burlingane, CA). For negative controls, we applied the same protocol to abomasum and small intestine sections but replaced the primary antibody with PBS. For positive controls, we used sections of jejunum and mesenteric lymph node from domestic cattle infected with *Haemonchus sp*. and *Cooperia sp*. No staining was detected in negative controls, and positive controls always yielded marked cytoplasmic staining of crypt enterocytes, plasma cells, and other leukocytes. IgA production in the mucosa was estimated based on the number of IgA-positive leukocytes in ten randomly selected microscope fields (400X) along the intestinal villi and crypts (48). We compared gut immunity between high and low FEC animals using data from control animals that did not receive anthelmintic treatment. For abomasal immunity, we used data from 14 animals (7 high FEC, 7 low FEC), and for intestinal immunity, we used data from 12 animals (6 high FEC, 6 low FEC).

### Genetics of worm resistance

To test for an underlying genetic basis to the worm resistance phenotypes characterized in our study population, we collected tissue samples from buffalo and stored them in silica gel at room temperature for up to 24 months prior to DNA extraction. DNA was extracted from tissue samples using the QIAGEN Blood & Tissue Kit (Valencia, CA) following the manufacturer’s protocol. DNA samples were genotyped at the BL4 microsatellite locus, which is located ∼3cM upstream of the interferon gamma gene (31). BL4 and other microsatellites in this chromsomal region have been associated with nematode resistance in sheep (28, 29, 49, 50). The BL4 locus has also previously been associated with nematode resistance in African buffalo (30). Genotyping was performed using PCR and fluorescently-labeled DNA fragment-visualization on an ABI3130xl automated capillary sequencer (Applied Biosystems) as described in (31). Allele sizes were determined using the ABI GS600LIZ ladder (Applied Biosystems). Chromatograms were analyzed by two independent technicians using GeneMapper v3.7. Tests for the presence of null alleles and deviations from Hardy-Weinberg proportions (HWP) showed no null alleles present at BL4 and no deviations from HWP (31).

### BTB infection and disease progression

We tested buffalo for BTB at each capture using a whole-blood interferon gamma (IFNγ) assay (BOVIGAM, Prionics, Switzerland) implemented according to the manufacturer’s instructions. We used bovine and avian tuberculin as stimulants for *in vitro* release of IFNγ in buffalo blood samples. BTB tests were considered negative if the optical density reading for bovine tuberculin-stimulated samples was below a cut-off of 0.375, and positive at readings of 0.375 or higher. Tests were also considered negative if the optical density reading for bovine tuberculin-stimulated samples failed to exceed the reading for avian tuberculin-stimulated samples by at least 10%, due to likely cross-reactivity to exposure to *M. avium*. This test protocol achieves a specificity of approximately 93.5% and a sensitivity of 85.4% (51). For each buffalo, we used a time series of 2-9 test results to classify BTB conversion status. For each animal, each test was interpreted in the context of the full time series of results to increase confidence in our assignment of BTB infection status (see (24) for details).

To quantify BTB progression, we performed gross and histopathological examinations on euthanized animals. First, animals were necropsied to assess the distribution of macroscopic BTB lesions in the lungs and respiratory lymph nodes (mandibular, parotid, palatine tonsil, retropharyngeal, caudal mediastenal, and tracheobronchial). Necropsies were performed by South African National Parks Veterinary Wildlife Services and South African State Veterinary Services veterinarians following standard protocols for BTB detection in buffalo (52). During necropsies, we serially sectioned the bronchial and retropharyngeal lymph nodes from every culled buffalo at 1-cm intervals to examine for tuberculous or other lesions, and immediately placed a 2 × 2 × 0.5 cm section from each lymph node (with or without lesions) in 10% neutral-buffered formalin (NBF). We thoroughly examined the lungs and diagrammed and measured all BTB lesions. We collected a 2 × 2 × 0.5 cm section from the edge of the largest pulmonary lesion of each animal into 10% NBF. From animals with no TB lesions, we collected a representative section of lung into 10% NBF. All formalin-fixed tissues were routinely processed for histopathology and stained with hematoxylin and eosin stains. Histological sections were examined by a board-certified veterinary pathologist who was blinded to the experimental groups. Sections of all tissues with tuberculous granulomas were examined for approximate percentage of the section affected by granulomatous inflammation and the percentage of the lesion that was necrotic. Lung sections were assessed for granuloma stage (53), and presence or absence of mineralization, fibrosis, and multinucleated giant cells. Retropharyngeal lymph node sections were assessed for stage (53), granuloma score (1 = focal, 2 = multifocal, 3 = coalescing, 4 = majority of the section), and presence or absence of mineralization, caseous necrosis, and multinucleated giant cells. We performed gross and histopathological examinations on 34 animals (18 high FEC, 16 low FEC; 21 treated, 13 control) that tested positive for BTB via BOVIGAM assay.

### Buffalo mortality

All study animals were fitted with radio-collars, so we continuously monitored the activity of all individuals to detect natural mortality events. For all possible mortality cases, we searched for the collars associated with these cases to determine the animal’s fate. Mortality was confirmed in 50 out of 67 cases and cause of death was ascertained where possible (e.g. predation). Three deaths that were deemed capture-related (e.g. a lion attack immediately after a captured animal was released) were excluded from the dataset. For each mortality event, the date of last recorded collar activity was assigned as the death date.

### Statistical analyses

#### Repeatability of fecal egg count and worm resistance phenotype

To examine the degree to which buffalo showed consistency in their level of worm egg shedding over time, we estimated the repeatability (R) of both continuous fecal egg count [log_10_ (FEC) + 1] and categorical worm egg shedding phenotype [high FEC or low FEC]. We estimated repeatability of FEC and worm phenotype using the LMM (gaussian) and GLMM (binary) methods, respectively, implemented with the *rptR* package (54) in R version 3.4.3. Confidence intervals were estimated using 1000 parametric bootstraps. For binary repeatability, we present R as a logit link-scale approximation, which provides a better estimate of the underlying propensity for an individual to express a high or low FEC. For both repeatability analyses, we restricted our analyses to the subset of control, non-anthelmintic treated buffalo (n = 103 individuals, 651 observations) in order to estimate the levels of natural consistency in worm shedding in the absence of anthelmintic treatment which we expected to disrupt egg shedding patterns.

#### Effect of initial worm phenotype on egg shedding over time and interaction with anthelmintic treatment

To evaluate whether worm phenotype (low *vs*. high) at an animal’s initial capture predicted worm egg shedding over the course of the study and how this related to anthelmintic treatment, we tested for an effect of worm phenotype at capture #1 on FEC at captures #2-*n* using a linear mixed model (LMM). Animal ID was included as a random effect in the model, and treatment status (control, treated), age, herd (CB, LS, Other), season (early dry, late dry, early wet, late wet), and capture interval (time since last capture) were included as fixed effects. We accounted for treatment, and an interaction between worm phenotype and treatment, since our past work showed that treatment significantly affected egg shedding rates (24). We also accounted for effects of age, herd, and season, since these factors are likely associated with differences in worm exposure and have a known influence on worm egg shedding in buffalo (24). Finally, we included capture interval to account for variable parasite re-accumulation in treated animals captured at different interval lengths. Prior to analysis, we normalized the distribution of the dependent variable (FEC) using log transformation. The model was run using the R packages *lme4* and *lmerTest*. Model validity was assessed by visual inspection of residuals as described in (55). The *difflsmeans* function in *lmerTest* was used to test for significant differences between least squared means. P-values were adjusted for multiple comparisons using Holm’s method implemented with the *p*.*adjust* function in the *stats* package.

#### Associations between worm resistance phenotype and gastrointestinal (GI) immunity and BL4 genotype

We tested for an association between worm resistance phenotype (high FEC vs. low FEC) and GI tract immune responses using Wilcoxon rank sum tests. This analysis focused only on non-anthelmintic-treated buffalo and we evaluated the effect of high vs. low status on numbers of eosinophils, mast cells, and IgA production in the abomasum and small intestine. We tested for associations between worm resistance phenotype and allelic variation at the BL4 locus (presence/absence of alleles 141 and 167) using chi-squared tests. We also used Wilcoxon rank sum tests to examine whether the presence of the two BL4 alleles were associated with differences in FEC at initial capture. For these genotype-based analyses, we included all sampled individuals (control and anthelmintic-treated) in the analyses because treatment should have had no impact on worm phenotype (which was classified at initial capture prior to first treatment) or genotype.

#### Survival analyses: effect of worm resistance phenotype on BTB infection probability and mortality

To examine the effects of worm resistance phenotype on BTB infection probability while accounting for anthelmintic treatment status, we added resistance status (high FEC vs. low FEC) as a predictor to a multivariate proportional hazards model previously used to test the effects of anthelmintic treatment on BTB infection risk (see (24)). The model included worm resistance phenotype, treatment, and herd of origin as predictors. BTB infection probability was estimated as the time from an animal’s initial capture to BTB infection or until the last BTB test. Including an interaction term between treatment status and worm resistance phenotype had no qualitative effect on the model results, so we do not report the interaction model. Our analysis included 195 individuals with known BTB infection histories that were also characterized for worm phenotype at initial capture. Data for individuals that did not test BTB positive by the end of the study period or that left the study due to death or emigration were right censored. The Efron method was used to address ties in event times, and the validity of the proportional hazards assumption was examined using Schoenfeld residuals (worm phenotype, low: χ^2^ = 0.403, *P* = 0.525, treatment, control: χ^2^ = 0.441, *P* = 0.521, herd, LS: χ^2^ = 0.005, *P* = 0.941, global model: χ^2^ = 0.936, *P* = 0.818). Cox regression models and residual checks were performed in R with the *survival* package. Hazard ratios, confidence intervals, and Wald tests for the model are reported in Table S3. The hazard ratio for the worm resistant phenotype variable (i.e. high FEC *vs*. low FEC) represents the instantaneous probability that a low FEC individual is more or less likely than a high FEC individual to acquire BTB.

We used a similar modeling procedure to test the effects of worm resistance phenotype on mortality risk for BTB-infected individuals. In this case, the multivariate model included worm resistance phenotype, treatment, herd, and age at first capture as predictors. A model including an interaction term between treatment status and worm resistance phenotype was undefinable due to low sample size. For all BTB-positives with known fates that were characterized for worm phenotype at initial capture (n = 56), we used the time from BTB conversion to either death or the end of the study as the response variable. Animals that did not die by the end of the study or those removed from the study for other reasons (e.g. emigration from the study area) were right censored. Four BTB-positive individuals that converted at their last capture were truncated from the dataset. Checks of the Schoenfeld residuals upheld assumptions of proportional hazards (worm phenotype, low: χ^2^ = 3.344, *P* = 0.0675, treatment, control: χ^2^ = 0.113, *P* = 0.7363, herd, LS: χ^2^ = 0.645, *P* = 0.4217, age: χ^2^ = 0.267, *P* = 0.6052, global model: χ^2^ = 6.307, *P* = 0.1774). Hazard ratios with confidence intervals and Wald tests are reported in the *SI Appendix* (Table S4).

To verify that the effect of worm resistance phenotype on buffalo mortality we observed was restricted to BTB-positive individuals and was not characteristic of BTB-negative individuals, we also examined the predictors of mortality risk in BTB-negative buffalo. There were 116 BTB-negative animals with known fates and worm phenotype characterizations. For these individuals we used the time from initial capture to death or the end of the study as the relevant response variable. Animals that did not die by the end of the study or those removed from the study for other reasons were right censored. The multivariate proportional hazards model had the same structure as the model run for BTB-positive individuals, and model results showed no effect of worm resistance phenotype on mortality risk (*SI Appendix*, Table S5). Residual checks upheld the assumptions of proportional hazards for this model (worm phenotype, low: χ^2^ = 0.085, *P* = 0.771, treatment, control: χ^2^ = 0.105, *P* = 0.706, herd, LS: χ^2^ = 0.702, *P* = 0.402, age: χ^2^ = 0.156, *P* = 0.693, global model: χ^2^ = 1.076, *P* = 0.898).

Finally, to assess whether our use of discretized FEC categories (i.e. worm resistance phenotypes) influenced conclusions about mortality patterns, we re-ran the mortality analyses described above for both BTB-infected and uninfected buffalo substituting continuous fecal egg count (FEC) for worm resistance phenotype. All other predictors remained the same and residual checks were performed as described. Model results are summarized in the *SI Appendix* (Tables S6-S7).

#### Effects of worm resistance phenotype and treatment on BTB progression

We examined the combined effects of worm resistance phenotype (high FEC vs. low FEC) and anthelmintic treatment on the progression of BTB disease using general and generalized linear models. Dependent variables were gross and histological measures of lung and lymph node pathology. Nominal variables were modeled using binomial GLM, ordinal variables were modeled using ordinal regression (R package: *ordinal*), and two continuous variables (# lung lesions and # positive lymph nodes) were modeled using negative binomial GLMs. For all models, worm resistance phenotype, treatment, duration of BTB infection, and the interaction between worm phenotype and treatment were included as predictor variables. A forward and backwards AIC-stepwise AIC-based model selection approach (implemented with the function *step*) was used to simplify each model and identify the key predictor variables explaining variation in disease progression.

## Author Contributions

VOE conceived the idea; VOE, SAB, GL, AEJ, and KS designed the study; VOE, SAB, PB, GL, MS, AEJ, and KS collected data; VOE analyzed the data and wrote the original manuscript; all authors provided feedback on the manuscript.

## Acknowledgments

We thank South African National Parks (SANParks) for permission to conduct this study in Kruger and the entire SANParks Veterinary Wildlife Services Department for invaluable assistance with animal captures and project logistics. Thanks to R. Spaan, J. Spaan, K. Thompson, B. Beechler, E. Gorsich, P. Snyder, J. Alagappan, S. Amish, L. Austin, L. Megow, K. Raum, N. Rogers and M. Smith for work on animal captures and sample processing. F. Quinn and P. Rohani provided valuable comments on the manuscript draft. Animal protocols for this study were approved by the University of Georgia and Oregon State University Institutional Animal Care and Use Committees (UGA AUP#: A2010 10-190-Y3-A5; OSU AUP#: 3822 and 4325).

## Funding

This study was supported by the National Science Foundation (NSF DEB-1102493) and National Institutes of Health (NIH 1R01GM131319) as part of the joint NSF/NIH/USDA Ecology of Infectious Diseases Grant Program.

## Competing interests

The authors declare no competing interests.

## Supplementary Information Appendix

**Figure S1.**
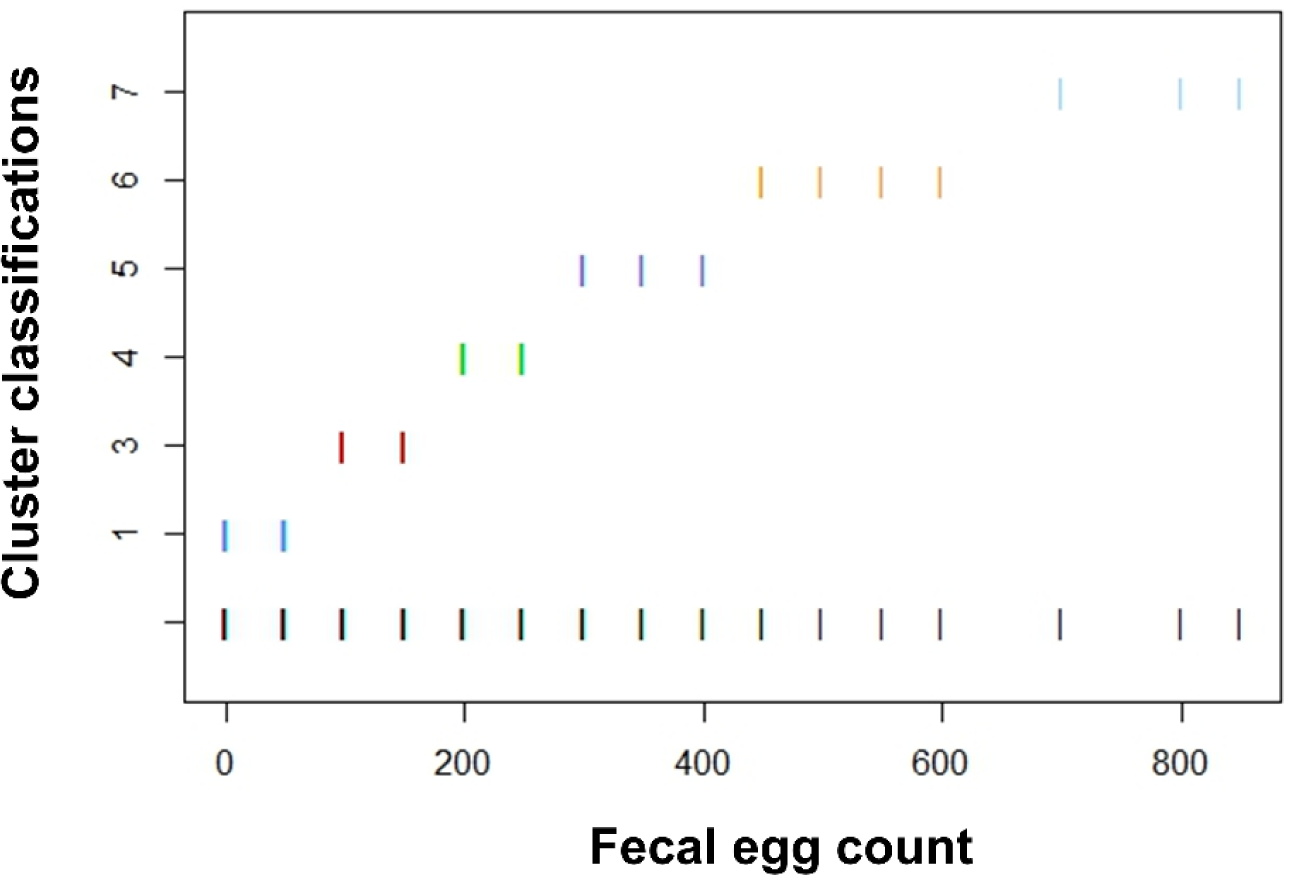
Fecal egg count (FEC) clusters identified from worm egg shedding patterns at initial sampling. Individual egg counts ranged from 0 to 850 eggs per gram of feces. An Expectation-Maximization clustering algorithm identified seven clusters in the data. Colored vertical lines on the graph indicate cluster assignments based on FEC value. Individuals shedding <100 worm eggs (Cluster 1) were assigned to the ‘low’ FEC category and individuals shedding ≥100 worm eggs (Clusters 2-7) were assigned to the ‘high’ FEC category.

**Figure S2.**
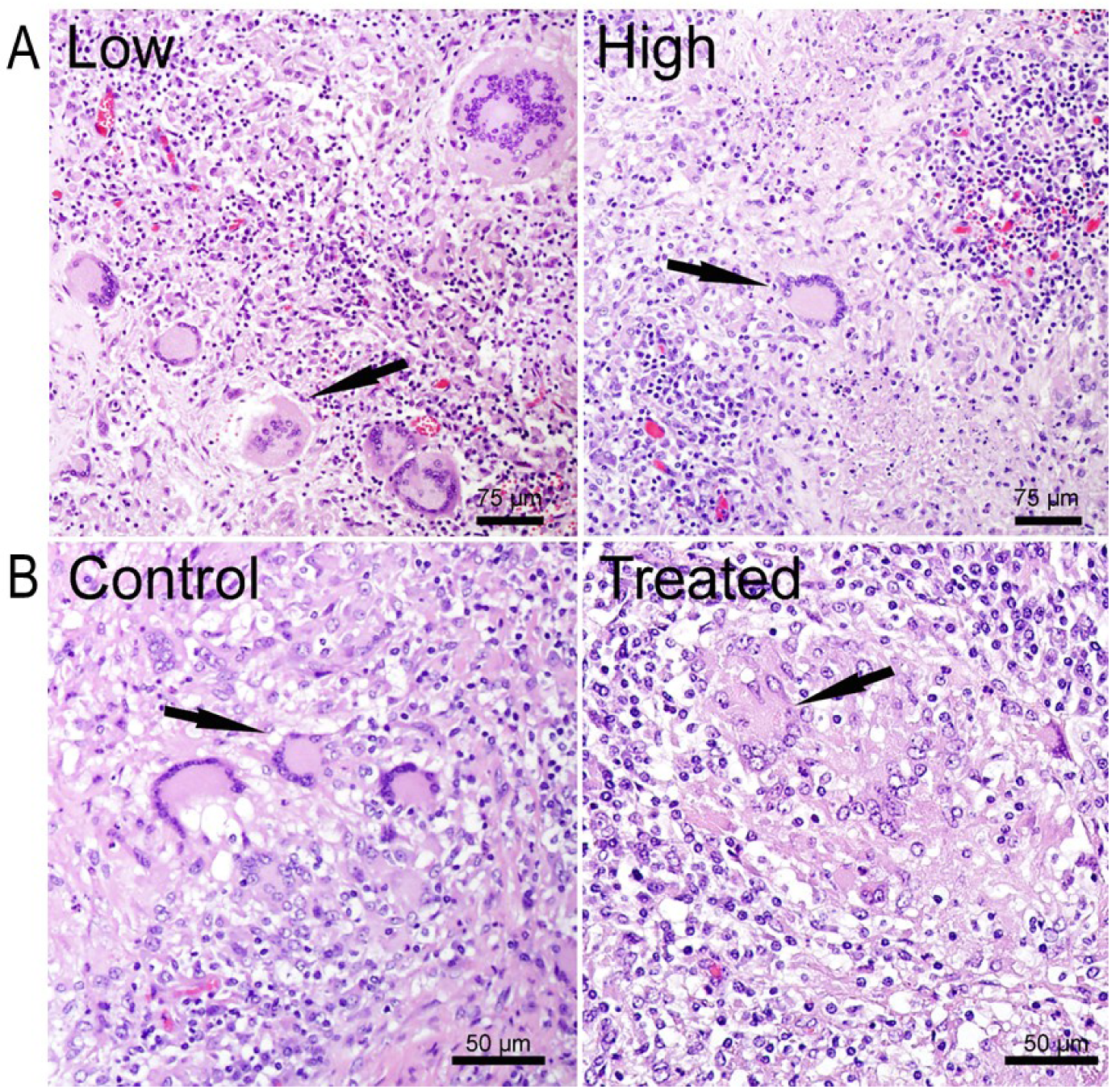
The effects of worm resistance phenotype and anthelmintic treatment on BTB progression were apparent in distinct tissues. BTB progressed more quickly in the lymph nodes of buffalo that were naturally resistant to worms and more slowly in the lungs of buffalo that received anthelmintic treatment. (A) Multinucleated giant cells (MNGCs, indicated by arrows) present within granulomatous lesions in the retropharyngeal lymph node of worm-resistant (low) and non-resistant (high) buffalo. Resistant individuals had significantly more MNGCs reflecting increased proliferation of mycobacteria and more severe disease. (B) MNGCs present within granulomatous lesions in the lungs of control and anthelmintic-treated buffalo. In this tissue, control individuals had significantly more MNGCs than treated individuals indicating increased bacterial proliferation and greater disease severity. For all **tables**, levels of significance are denoted by asterisks (***p <0.001, **p<0.01, *p<0.05).

**Table S1.** Worm phenotype (high or low) at initial capture, anthelmintic treatment, and the interaction between both factors as predictors of worm egg shedding over the entire study duration (n = 209 animals, 1101 observations). Linear mixed model (LMM) with Animal ID included as a random effect; Type III ANOVA table.

|  | df | F value | P |
| --- | --- | --- | --- |
| Worm phenotype | 1 | 31.3 | <0.001*** |
| Treatment | 1 | 61.5 | <0.001*** |
| Season | 3 | 5.2 | 0.0145* |
| Herd | 2 | 2.22 | 0.109 |
| Age | 1 | 2.88 | 0.09 |
| Capture interval | 1 | 29.69 | <0.001*** |
| Worm phenotype × Treatment | 1 | 11.02 | <0.0011** |

**Table S2.** Least squares means for worm phenotype, anthelmintic treatment, and the interaction between both factors (from Table S1), showing the magnitude of differences in worm egg shedding between groups. Significance levels were adjusted for multiple comparisons using Holm’s method.

|  | Estimate | Std. Error | df | t value | P |
| --- | --- | --- | --- | --- | --- |
| High – Low | 0.499792 | 0.089274 | 208.2 | 5.5984 | <0.001*** |
| Treatment – Control | -0.686992 | 0.087596 | 201.7 | -7.8427 | <0.001*** |
| Treated High – Treated Low | 0.209664 | 0.125599 | 208 | 1.6693 | 0.193 |
| Treated High – Control High | -0.97712 | 0.136621 | 205.7 | -7.1521 | <0.001*** |
| Treated High – Control Low | -0.1872 | 0.126564 | 209.6 | -1.4791 | 0.193 |
| Treated Low – Control High | -1.186785 | 0.123562 | 200.3 | -9.6048 | <0.001*** |
| Treated Low – Control Low | -0.396864 | 0.109348 | 193.4 | -3.6294 | 0.00109** |
| Control High – Control Low | 0.789921 | 0.12426 | 200 | 6.357 | <0.001*** |

**Table S3.** Proportional hazards model of BTB-infection risk including worm resistance phenotype, anthelmintic treatment status, and herd as predictors.

|  | **Hazard ratio** | **95% CI** | **Z** | **P** |
| --- | --- | --- | --- | --- |
| Resistance phenotype (low vs. high) | 0.708 | 0.427-1.177 | -1.331 | 0.183 |
| Treatment (control vs. treated) | 0.965 | 0.592-1.572 | -0.145 | 0.885 |
| Herd | 1.499 | 0.897-2.504 | 1.546 | 0.122 |

Whole model: Likelihood ratio test= 3.36, 3 df, $P = 0.339$ , $n = 195$

**Table S4.** Proportional hazards model of mortality risk in BTB-positive individuals including worm resistance phenotype, anthelmintic treatment status, herd, and age at first capture as predictors.

|  | **Hazard ratio** | **95% CI** | **Z** | **P** |
| --- | --- | --- | --- | --- |
| Resistance phenotype (low vs. high) | 5.6758 | 1.079-29.8562 | 2.05 | 0.04039* |
| Treatment (control vs. treated) | 10.5482 | 1.966-56.5836 | 2.749 | 0.00598** |
| Herd | 0.2548 | 0.07318-0.8872 | -2.148 | 0.03171* |
| Age | 1.0391 | 1.0056-1.0737 | 2.297 | 0.02162* |

Whole model: Likelihood ratio test= 19.17, 4 df, $P = 0.0007283$ , $n = 56$

**Table S5.** Proportional hazards model of mortality risk in BTB-negative individuals including worm resistance phenotype, anthelmintic treatment status, herd, and age at first capture as predictors.

|  | **Hazard ratio** | **95% CI** | **Z** | **P** |
| --- | --- | --- | --- | --- |
| Worm resistance phenotype<br>(low vs. high) | 0.7739 | 0.3795-1.578 | -0.705 | 0.4809 |
| Treatment<br>(control vs. treated) | 0.5596 | 0.2676-1.17 | -1.542 | 0.123 |
| Herd | 0.4182 | 0.1833-0.954 | -2.072 | 0.0383* |
| Age | 0.976 | 0.952-1.001 | -1.914 | 0.0556 |

Whole model: Likelihood ratio test = 13.19, 4 df, $P = 0.01038$ , $n = 116$

**Table S6.** Proportional hazards model of mortality risk in BTB-positive individuals using continuous fecal egg count (FEC) rather than discretized resistance phenotype (low *vs*. high) as a response variable. Additional predictors include anthelmintic treatment status, herd, and age at first capture.

|  | **Hazard ratio** | **95% CI** | **Z** | **P** |
| --- | --- | --- | --- | --- |
| FEC | 0.9944 | 0.98835-1.0005 | -1.812 | 0.07003 |
| Treatment<br>(control vs. treated) | 15.5481 | 2.60420-92.8278 | 3.010 | 0.00261** |
| Herd | 0.2360 | 0.06167-0.9029 | -2.109 | 0.03494* |
| Age | 1.0470 | 1.01171-1.0835 | 2.625 | 0.00865** |

Whole model: Likelihood ratio test= 18.46, 4 df, $P = 0.001005$ , $n = 56$

**Table S7.** Proportional hazards model of mortality risk in BTB-negative individuals using continuous fecal egg count (FEC) rather than discretized resistance phenotype (low *vs*. high) as a response variable. Additional predictors include anthelmintic treatment status, herd, and age at first capture.

|  | **Hazard ratio** | **95% CI** | **Z** | **P** |
| --- | --- | --- | --- | --- |
| FEC | 0.998677 | 0.9954-1.0019 | -0.799 | 0.4241 |
| Treatment<br>(control vs. treated) | 0.613188 | 0.2902-1.2957 | -1.281 | 0.2001 |
| Herd | 0.371645 | 0.1624-0.8504 | -2.344 | 0.0191* |
| Age | 0.974416 | 0.9507-0.9987 | -2.064 | 0.0390* |

Whole model: Likelihood ratio test = 13.44, 4 df, $P = 0.009295$ , $n = 116$

## Notes

### Competing Interest Statement

The authors have declared no competing interest.

